# Simpler protein domain identification using spectral clustering

**DOI:** 10.1101/2024.02.10.579762

**Authors:** Frédéric Cazals, Jules Herrmann, Edoardo Sarti

## Abstract

The decomposition of a biomolecular complex into domains is an important step to investigate biological functions and ease structure determination. A successful approach to do so is the `SPECTRUS` algorithm, which provides a segmentation based on spectral clustering applied to a graph coding interatomic fluctuations derived from an elastic network model.

We present `SPECTRALDOM`, which makes three straightforward and useful additions to `SPECTRUS`. For single structures, we show that high quality partitionings can be obtained from a graph Laplacian derived from pairwise interactions–without normal modes. For sets of homologous structures, we introduce a Multiple Sequence Alignment mode, exploiting both the sequence based information (MSA) and the geometric information embodied in experimental structures. Finally, we propose to analyse the clusters/- domains delivered using the so-called *D*-family-matching algorithm, which establishes a correspondence between domains yielded by two decompositions, and can be used to handle fragmentation issues.

Our domains compare favorably to those of the original `SPECTRUS`, and those of the deep learning based method `Chainsaw`. Using two complex cases, we show in particular that `SPECTRALDOM` is the only method handling complex conformational changes involving several sub-domains. Finally, a comparison of `SPECTRALDOM` and `Chainsaw` on the manually curated domain classification `ECOD` as a reference shows that high quality domains are obtained without using any evolutionary related piece of information.

`SPECTRALDOM` is provided in the Structural Bioinformatics Library, see http://sbl.inria.fr and https://sbl.inria.fr/doc/Spectral_domain_explorer-user-manual.html.

## 1 Introduction

### 1.1 Protein segmentation and dynamic domains

#### Molecular functions, motions, and domains

Molecular motions within proteins span *∼* 15 orders of magnitude [1], and their prediction is key to shed light on complex biological functions. A classical strategy to study these complex phenomena consists of abstracting a molecular system in a hierarchical fashion, with rigid domains connected by linkers. While this decomposition may be carried out in a recursive fashion, the simplest model consists of a flat hierarchy–rigid domains connected by linkers. Such a model corresponds to a segmentation such that interactions within domains are tight, while those across domains are looser. Identifying domains based on such interactions requires taking into account the atomic packing, the presence of disulfide bonds, hydrogen bonding, and naturally atomic movements. In modeling a biomolecule as a graph whose units are amino acids or bases, with an edges between interacting units, the previous physical intuition calls for methods reminiscent of graph clustering and/or community detection. We now briefly review two classes of methods for this task.

#### Domains via (non) supervised approaches

Spectral graph partitioning techniques form a rich class of methods which may be interpreted in terms of graph min cuts as well as random walks and Markov chains [2]. The domain partitioning algorithm `SPECTRUS` [3] falls in this vein, as it relies on spectral clustering based on a similarity matrix derived from fluctuations and normal modes–details thereafter. A similar three step partitioning procedure is `SWORD2` [4]. The first step performs an assignment of secondary structure elements (SSE) with eight conformational states. The second one is based on a contact map defined from a logistic function applied to inter-residue distances this step identifies protein units (PUs) using an iterative hierarchical clustering of the contact probability map. The final step reconstructs domains by gradually merging PUs, maximizing the compactness of domains and their separation. The algorithm `UniDoc` defines domains by maximizing intra-domain interactions and minimizing inter-domain interactions. Inter/intra domain scores are defined from a logistic function on the pairwise *C*_*β*_ (*C*_*α*_ for glycine) distances [5]. In a first step, the protein is split into fragments (using so-cal. led continuous and discontinuous splits) based on a graph min cut. The second one iteratively merges fragments into domain to maximize the interaction inside each domain.

Supervised approaches were also recently developed. The method `FUPred` [6] couples contact maps yielded by deep residual neural networks and coevolutionary precision matrices. `Chainsaw` [7] uses a residual convolutional neural network to predict the aforementioned adjacency matrix over residues. The structure is converted into five feature channels (pairwise *C*_*α*_ distances, and four channels for predicted SSE (helix vs strand, within vs boundary of SSE)). The features are converted into a pairwise probability matrix using a deep convolutional network initially designed for structure prediction–a variant of `trRosetta` [8]. The learning process minimizes the cross entropy between the predicted soft adjacency matrix and the one representing the probability of residue co-occurrence in the same domain.

The Domain Parser for Alphafold models (DPAM) targets `AlphaFold2`models [9]. This raises two new challenges: the large number of structures, and the requirement to handle non domain regions which are unsuitable for domain classification (IDR, linkers, coiled coil, etc). Of particular interest in this context is the Predicted Aligned Error (PAE) provided by `AlphaFold2`. A *combined probability* for pairs of residues is defined, based on the PAE, the inter-residue distance, the HHsuite support, and the Dali support. The weights are learned to match a library of known domains. Using a partitioning of the protein into residues segments of length five, an iterative clustering of residues is performed to obtain domains.

### 1.2 Contributions

We present three straightforward and useful simplifications and extensions to `SPECTRUS`. First, for single structures, we show that high quality partitioning can be obtained from a graph Laplacian derived from pairwise interactions of several types (covalent, non-covalent, hydrogen bonds, salt-bridges, etc). This mode, termed the *diffusion map* (DM) mode is a significant simplification of `SPECTRUS` algorithm [3], which derives atomic fluctuations from the normal modes of an elastic network model (ENM) [10]. The DM mode indeed relies on a single eigenproblem (used in spectral clustering) instead of two (normal modes, spectral clustering). Denoting *n* the number of residues, the fact that the latter operates on a matrix of size *n* × *n* instead of 3*n* × 3*n* (for normal modes) yields a speedup of at least one order of magnitude–considering a quadratic lower bound for sparse eigenproblems.

Second, for sets of structures, we introduce a multiple sequence alignment (MSA) mode, in which fluctuations are derived from valid positions within the MSA. Using various structural measures (lRMSD from flexible structural alignments and combined RMSDs), we show in particular that the domains obtained are useful to study conformational changes and molecular motions.

Third, we propose to analyse the clusters/domains delivered in a hierarchical fashion, using an edit distance between clusters, and we use this technique to reduce domain fragmentation issues faced by most if not all previous methods.

We show that our domains compare favorably to those output by two reference methods, the original `SPECTRUS`, and those of the deep learning based method `Chainsaw`. We show in addition that our clusters are more coherent with those from the reference database `ECOD`, despite the fact that we do not use any evolutionary related piece of information.

The software `SPECTRALDOM` is made available in the Structural Bioinformatics Library, see http://sbl.inria.fr and https://sbl.inria.fr/doc/Spectral_domain_explorer-user-manual.html.

## 2 Methods: SPECTRUS and SPECTRALDOM

### 2.1 The original SPECTRUS algorithm

For the sake of completeness and exposure, let us review briefly the e `SPECTRUS` algorithm [3]. The method relies on the following intuition to define domains: atomic fluctuations within (resp. across) domains are tame (resp.large). For a predefined number of clusters/domains *k*, the algorithm is as follows:

1. Compute a fluctuation matrix assessing the distance variation between any two residues in the chain. Denoting *d*_*ij*_ the distance between the *C*_*α*_ carbons of amino acids *i* and *j*, the quantity of interest is the standard deviation

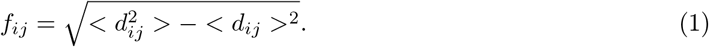

For a single structure [3], fluctuations are derived from from normal modes associated with a harmonic model [10]. For several homologous chains, one estimates Eq. (1) from the distances observed in the conformations processed.
2. Convert the fluctuations *f*_*ij*_ into similarities *w*_*ij*_ via thresholding and exponentiation. That is, with *σ* a conservative measure for intra-cluster/domain dissimilarity and *d*_max_(= 10) a distance threshold:

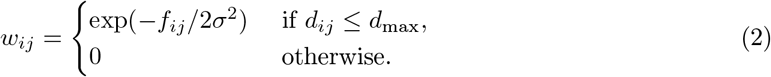

This results in a weighted graph whose vertices are the individual residues. Intuitively, the weight *w*_*ij*_ on edge *i* to *j* can be seen as the probability for residues *i* and *j* to be in the same cluster/domain.
3. Apply spectral clustering to the previous graph [11]. This step involves clustering vectors on the unit sphere *S*^*k*−1^, which is done with *k*-medoids in `SPECTRUS`.

To identify suitable values of *k*, the previous steps are repeated. A classical way to assess the significance of a clustering is to resort to a null model [12, 13]. For `SPECTRUS`, the quality of the domains obtained for each *k* is measured by a score (Score_*k,n*_, Eq. (4)) based on a ratio involving a null model consisting of random points on *S*^*k*−1^. See details in Sec. S1. The boxplot of this score as a function of *k* is termed the *quality plot*. The values of *k* interest are the local maxima of the quality plot with low variance of the score Score_*k,n*_. In increasing *k*, one expects this variance to grow: a larger *k* entails more partitions, some with less pertinent boundaries, whence a larger score and variance. As we shall see, this behavior is indeed observed in practice and is useful in picking appropriate values of *k*.

### 2.2 SPECTRALDOM

We now present our three additions to `SPECTRUS`.

#### Direct segmentation of a single model: the diffusion map (DM) mode

The original `SPECTRUS` for single structures uses spectral clustering based on similarities defined from the normal modes of an elastic network model (ENM). A more direct route consists of defining the weights *w*_*ij*_ directly from the spring constants of the ENM, thus removing the normal modes calculation. Consider three stiffness constants 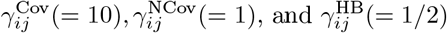, for covalent, non covalent, and hydrogen bonds respectively. (We note in passing that additional terms *e*.*g*. for salt bridges can easily be accommodated to weight contacts.)

Because non covalent contacts are detected using a distance threshold, denoting *d*_*ij*_ the distance between the *C*_*α*_ carbons of two interacting residues, we soften the stiffnesses *γ*_*ij*_ as follows:

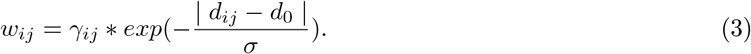

We take *d*_0_ = 4.5Å as typical distance between consecutive *C*_*α*_s along the backbone, and *σ* = 1*/*2.

Note that two amino acids may interact in different ways. For example, two a.a. on the same generatrix of a *α*-helix interact non-covalently, but also via hydrogen bonding. We therefore define two modes to define the weight *w*_*ij*_ of two residues with multiple interaction types, the `Weight-Max` (resp. `Weight-Sum`) mode in which we take the max (resp. the sum) of individual stiffnesses. In our helix example, one gets *γ*_*ij*_ = max(*γ*^NCov^, *γ*^HB^), and *γ*_*ij*_ = *γ*^NCov^ + *γ*^HB^ respectively. The resulting weights are directly passed to spectral clustering based on the symmetric Laplacian [11].

This diffusion map (DM) mode preserves the anisotropy of fluctuations embodied in normal modes (if any) via the anisotropy of contacts (if any). But it has two direct advantages over the ENM mode (original `SPECTRUS`): (i) the segmentation requires contacts/distances only rather than coordinates used to derive the Hessian of the ENM, (ii) the matrix diagonalization operates on a sparse matrix of size *n* × *n* rather than 3*n* × 3*n*.

##### Remark 1

*In processing PDB files, hydrogen bonding information is retrieved from secondary structures. A mode loading a* DSSP *file is available, but was not used for experiments*.

#### SPECTRUS for multiple homologous structures: the MSA mode

Consider multiple structures for which a MSA is available. We cover the case of conformations of the same protein using this model also, since experimental structures of the same protein often harbors addition/deletion at the N-tear and/or C-ter of the chain. A position in the MSA is termed *valid* provided that (i) the corresponding chains have no gap at this position in the MSA, and (ii) the corresponding residues have been solved in the individual experimental structures. The fluctuation of Eq. (1) can now be computed for each pair of valid indices in the MSA, and used for spectral clustering again. One obtains a partitioning of the structures into domains which are *compact/rigid*, but move with respect to one another.

While conceptually straightforward, the MSA mode requires a careful treatment of all indices involved (resids in structures which are not contiguous in general, sequence indices, alignment indices). See Sec. S1.2.

##### Remark 2

*The ability to handle homologous structures via a MSA deserves a comment in terms the sparsity of alignments and structures handled. If too many residues are invalid (Section S1, Defs. 2 and 3), the similarity matrix is too sparse, which results in a large number of clusters. Intuitively, one should recall that the eigenspace associated to the value 0 of the graph Laplacian encodes connected components of the graph*.

#### Fragmentation issues

The spectral decomposition may yield fragmentation issues, when a region (*e*.*g*. a region from an helix) is assigned to a nearby domain. For `SPECTRUS` or `SPECTRALDOM`, this happens when the distance *d*_*ij*_ between two residues is small, which results in large weight *w*_*ij*_ – Eq. (3).

We handle such issues using an algorithm identifying the correspondences between the domains of two candidate decompositions, called the *D*-family-matching algorithm [14]. Let 𝒞_1_ = {*C*_1,*i*_} and 𝒞_2_ = {*C*_2,*j*_} be two decompositions into domains associated to two values *k*_1_ and *k*_2_. In short, the *D*-family-matching creates groups of clusters (domains in our case) within 𝒞_1_ and likewise within 𝒞_2_, called *meta-clusters*, with a one-to-one correspondence between these meta-clusters. The method relies on the complete edge-weighted bipartite graph 𝒞_1_ × 𝒞_2_, the weight of edge (*C*_1,*i*_, *C*_2,*j*_) being equal to the number of points (amino-acids in our case) shared by *C*_1,*i*_ and *C*_2,*j*_. It is governed by a scale parameter *D* counting the maximum number of steps (zigzags) one can make to join a cluster from 𝒞_1_ to a cluster of 𝒞_2_–formally the diameter of the subgraph induced by meta-clusters. The number of meta-clusters obtained is dictated by the combinatorial optimization problem solved. See [14] for details, as well as the software in the Structural Bioinformatics Library https://sbl.inria.fr/doc/D_family_matching-user-manual.html. Consider a chain which faces a fragmentation issue (Figs. S2 and S3). Let us consider two decompositions 𝒞_1_ and 𝒞_2_ of this chain into domains. Further decompose each domain *C*_1,*i*_ ∈𝒞_1_ (and likewise *C*_2,*j*_ ∈𝒞_2_) into contiguous stretches along the sequence of the chain, yielding *C*_1,*i*_ = ∪ _*j*_*s*_1,*i,j*_ – with *i* (resp. *j*) the index of the domain (resp. stretch). To possibly re-assign a mis-assigned stretch *s*_1,*i,j*_ to the correct domain of 𝒞_1_, we proceed in three steps. (NB: the method is mutatis mutandis for a stretch from a domain from 𝒞_2_.) First, using *D*-family-matching with *D* = 2, we compute meta-clusters *M*_*k*_, *k* = 1, …, *K* to match groups of {*C*_1,*i*_} with groups of {*C*_2,*j*_}. The clusters of 𝒞_1_ (resp. 𝒞_2_) aggregated by *M*_*k*_ are denoted 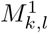 (resp. 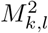). Second, we compute the intersection size 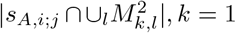, *k* = 1, …, *K*. Let *o* ∈ 1, …, *K* be the index of the meta-cluster yielding the best score for *s*_1,*i,j*_. Consider the set 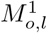 of clusters/domains from 𝒞_1_ associated with this meta-cluster *M*_*o*_. Because *s*_1,*i,j*_ is contiguous along the sequence, it is flanked by exactly one or two domains from 𝒞_1_. Let the *flanking number* the number of such domains from 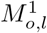. If the flanking number is one, we assign *s*_1,*i,j*_ to that domain. If not, we do not perform any update: a value of zero would mean that the stretch is not contiguous to a domain from 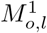; and a value *>* 1 would trigger the merge of two domains identified by `SPECTRALDOM`.

##### Remark 3 (Clustering on *S*^*k*−1^ and quality plots.)

*Spectral clustering requires clustering unit vectors on the unit sphere S*^*k*−1^. *We use* `k-means++` *instead of* `k-medoids` *in* `SPECTRUS` *[15]. Due to the hardness of* `k-means` *clustering and the random initialization of seeds in* `k-means++`, *several clusterings are in general obtained for a given value of k. Whence the boxplot associated to each value of k on our quality plots. In practice, the default range of values explored by* `SPECTRALDOM` *is* [*k*_*min*_ = 2, *k*_*max*_ = 10]. *On the quality plots, the values of k such that primary sequence size divided by the number of domains is less than 35 amino acids (default size for a helix) are shaded, as decomposing a structure into SSE is not the goal pursued*.

### 2.3 Implementation

The implementation is provided in the Structural Bioinformatics Library [16]. See Sec. S1 (Algo. 1) for the pseudo-code, and https://sbl.inria.fr/doc/Spectral_domain_explorer-user-manual.html.

## 3 Experiments

We present three complementary experiments: Sec. 3.1 illustrates the ability of `SPECTRALDOM` to handle challenging case for competing methods; Sec. 3.2 compares the MSA and DM modes of `SPECTRALDOM` to study conformational changes and molecular motions; Sec. 3.3 studies the coherence between the domains found by `SPECTRALDOM` and those of the `ECOD` database [17].

Experiments in the DM mode for single structures use the `Weight-Sum` mode for weights–Sec. 2.2, and exploit hydrogen bonding information from SSE elements in PDB file headers. Default values were used for the stiffness constants, that is 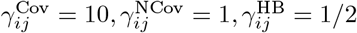 The distance threshold used to consider the fluctuation of two residues was wet to *d*_max_(= 10) to avoid too sparse graphs favoring an over-segmentation.

### 3.1 Comparisons between SPECTRUS, Chainsaw, and SPECTRALDOM

#### Rationale: assessing partitioning and fragmentation

In order to illustrate the efficacy of `SPECTRALDOM`, we present in this section three examples of increasing difficulty: the first is extracted from the `SPECTRUS` test set, the second from the `Chainsaw` test set, and the third is a system we recently studied and for which we have clear dynamical information. The complete test sets from the `SPECTRUS` and `Chainsaw` works are included in the SBL package.

#### Results: Escherichia coli adenylate kinase

The Escherichia coli adenylate kinase (PDBid 1ake and 4ake), studied in [3], is a monomeric enzyme of about 200 amino acids, which freely interconverts between open and close conformations (Fig. 1). When `SPECTRUS` is given the two conformations, it finds the correct number of clusters *k* = 3, where two clusters correspond to the ATP binding domain (LID) and the AMP binding domain (NMP) whereas the third includes the rest of the structure. This decomposition incurs two spurious mis-assignment due to fragmentation issues (Fig. 1, `SPECTRUS`, circled areas). `SPECTRALDOM` in its MSA mode reproduces this result, but does not suffer from these mis-assignments. `SPECTRALDOM` also finds the same partition from the sole open conformation (PDBid 4ake), which confirms the robustness of the method. On the same system, `Chainsaw` was not able to detect multiple domains.

**Figure 1:**
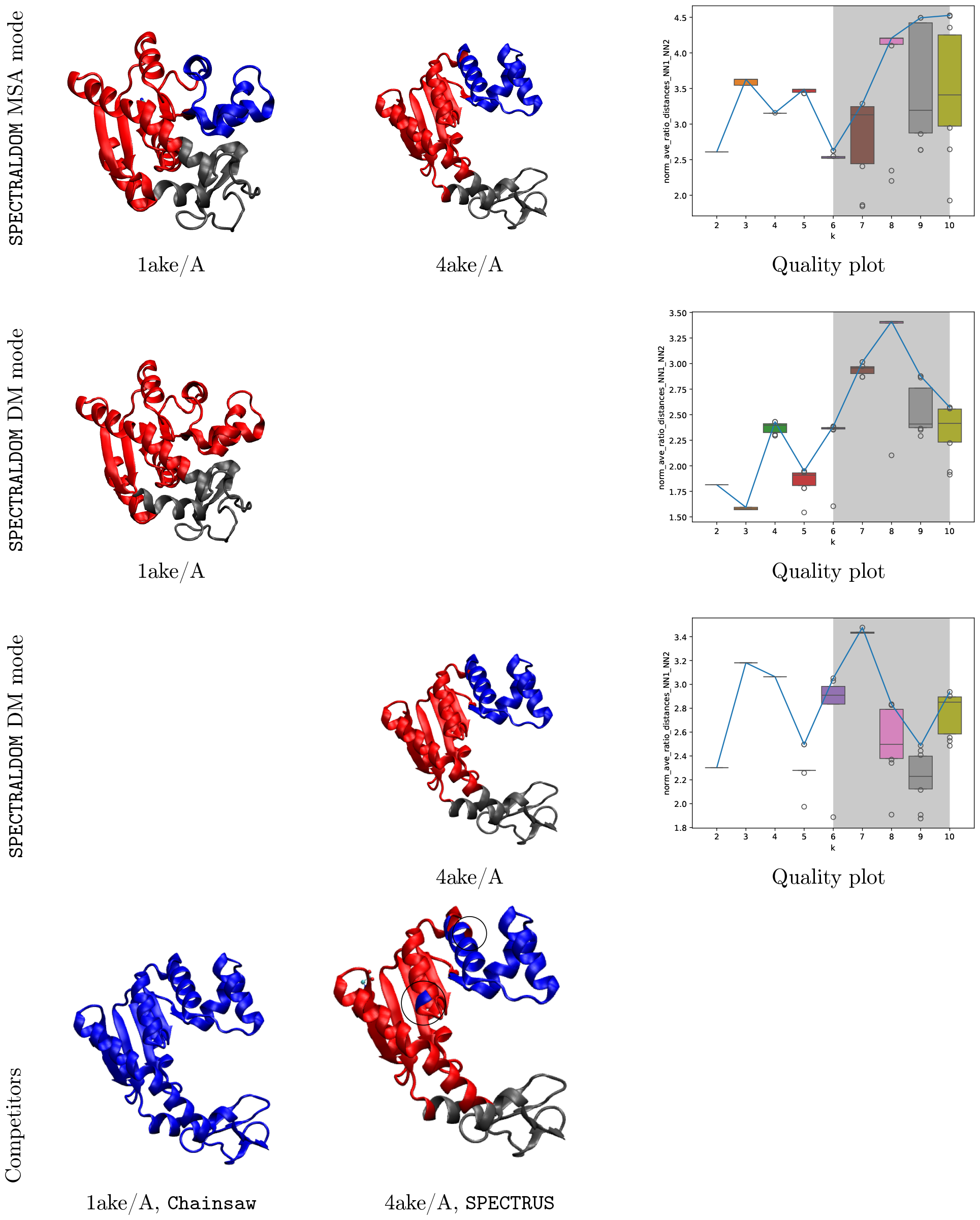
Escherichia coli adenylate kinase: comparison of the SPECTRALDOM modes MSA and DM. close and open conformations (PDBid 1ake, 4ake). Crystal structures: closed: 1ake/A; open: 4ake/A. (Row 1) MSA mode applied to chains A of 1ake and 4ake. (Row 2) DM mode for 1ake/A. (Row 3) DM mode for 4ake/A. (Row 4) Comparison with `SPECTRUS` and `Chainsaw`

#### Results: human serum transferrin

The human serum transferrin (PDBid 1a8e) presents two domains one of which is discontinuous (i.e. formed by the union of two continuous polypeptide segments). This poses a challenge for algorithms that are intrinsically based on sequence such as `Chainsaw`, but does not pose additional complexity for graph-based approaches such as `SPECTRALDOM`. `Chainsaw` and `SPECTRALDOM` both correctly predict the partition (Fig. 2), but `Chainsaw` incorrectly assigns the last residue of the C-terminal helix. `SPECTRUS` also correctly finds *k* = 2, but misassigns half of the same C-terminal helix. Whereas it is disputable whether the terminal helix must belong to one domain or the other, we find the partition of `SPECTRALDOM` more coherent than the others.

**Figure 2:**
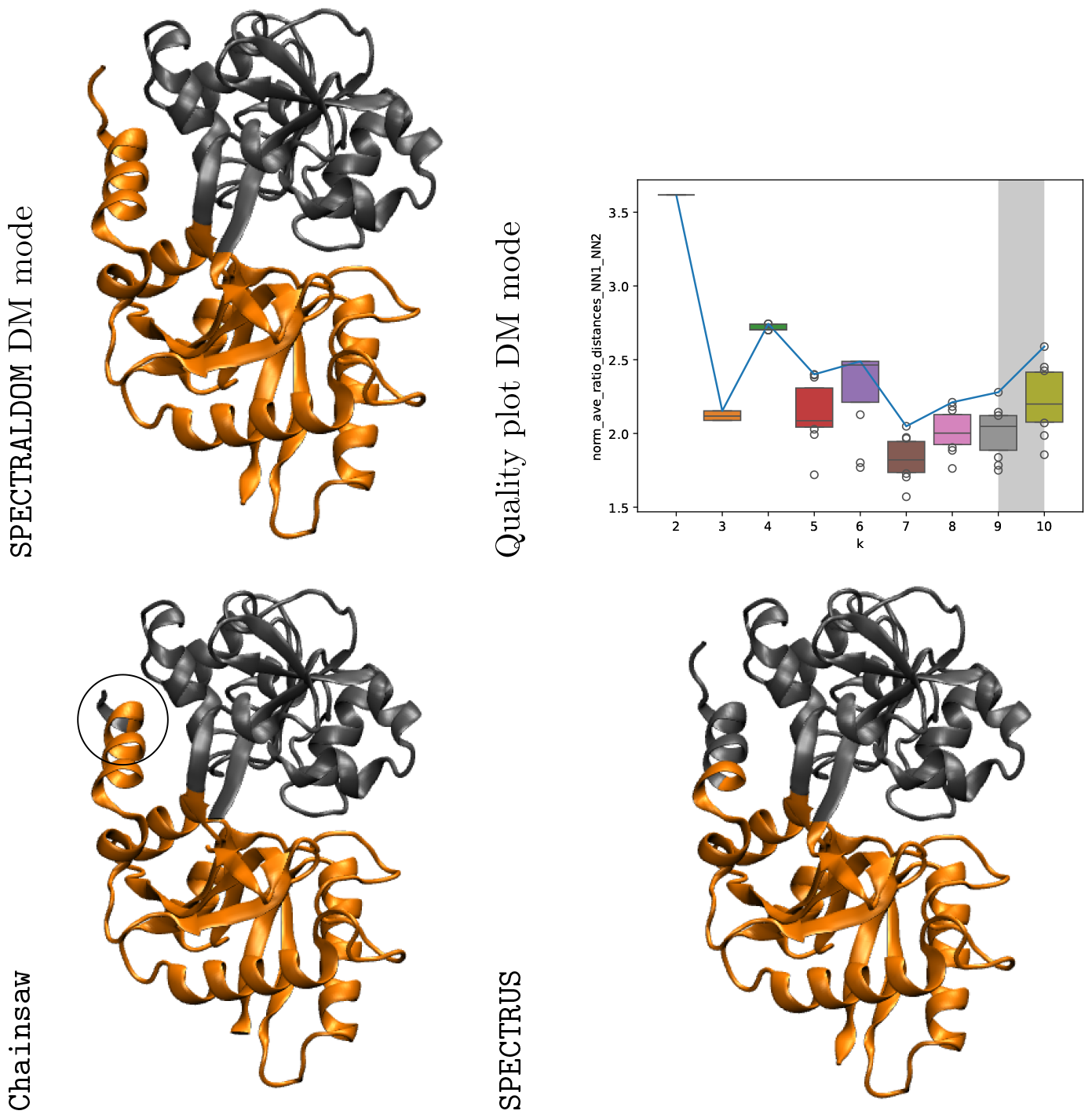
Human serum transferrin (PDBid 1a8e/A), DM mode. The protein has two dynamical domains one of which composed by non-contiguous segments. `SPECTRALDOM`, `Chainsaw`, and `SPECTRUS` recover the correct number of clusters, with slight differences in the partition on the terminal helix.

#### Results: acriflavine resistance B

Acriflavine resistance B (AcrB, PDBid 4zit) is a transmembrane drug/proton antiporter serving as drug efflux pump in Gram-negative bacteria. It is a homotrimer where each monomer cycles between three consecutive states that compose the alternating access mechanism. These conformational changes rely on the interplay of 7 quasi-rigid dynamical domains and 1 disordered domain. `SPECTRALDOM` finds a partition in 4 clusters that are unions of the real quasi-rigid domains (Fig. 3). So does `Chainsaw` (although its partition is not symmetric) and `SPECTRUS` (although its domains include several incorrect fragments). `SPECTRALDOM` is the only one to suggest the correct partition as second-best candidate.

**Figure 3:**
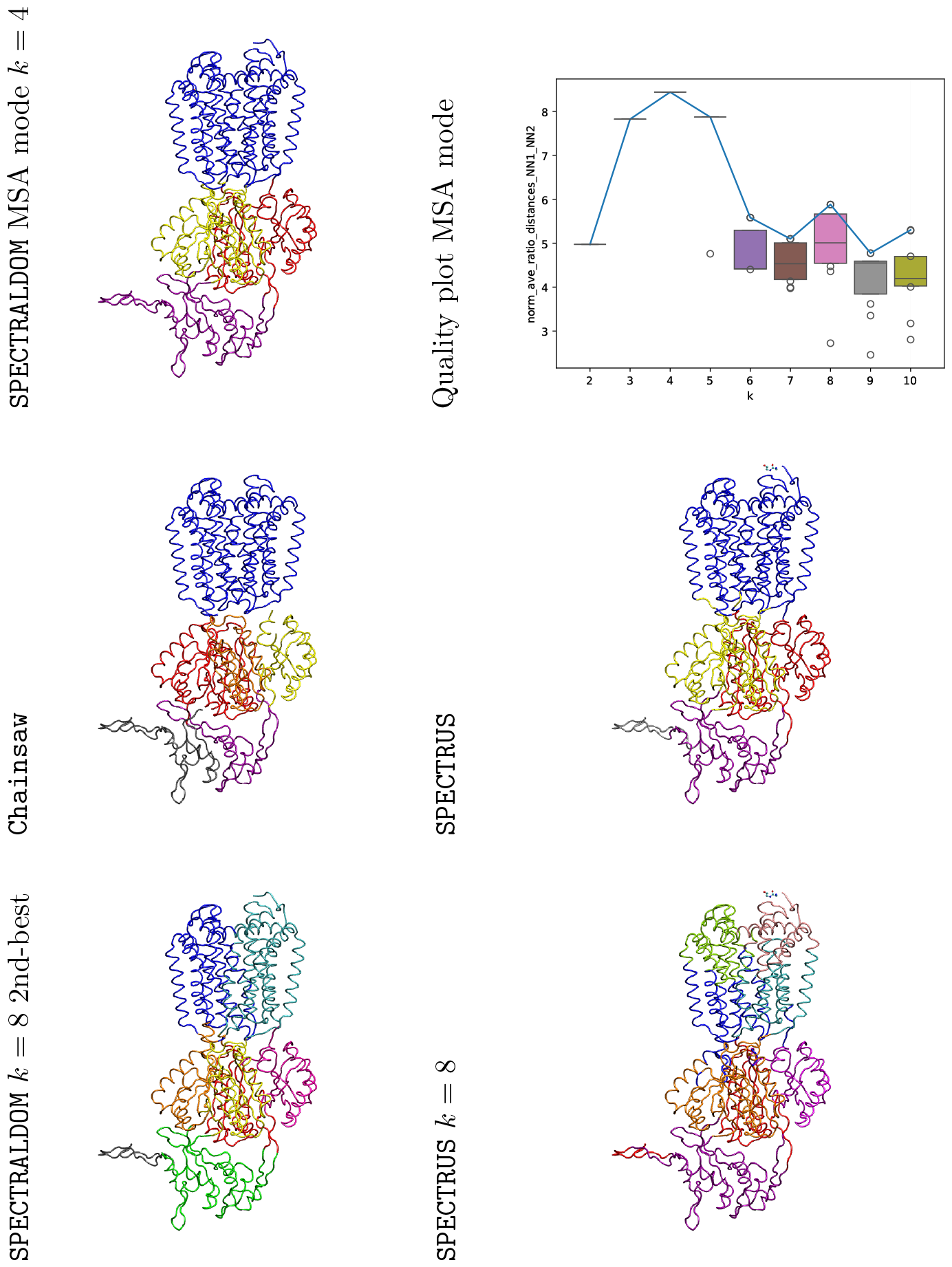
Monomer of the trimeric Acriflavine resistance B (AcrB, PDBid 4zit/A), MSA mode. The monomer can be decomposed in 7 quasi-rigid domains plus 1 disordered domain. `SPECTRALDOM` is the only algorithm to suggest the correct repartition as second-best choice, the best one still being correct yet more coarse-grained.

#### Remark: quality score variance as a function of

*k* It is interesting to note that in all three examples, the quality score variance increases with increasing *k*. Although qualitative, this trend reveals that finer partitions are on average less stable than coarser ones – larger values of *k* increase the number of possible partitions, some of which incur a large variance.

### 3.2 Domains: prediction in case of conformational changes

#### Rationale: modeling conformational changes

We wish to compare the DM and MSA modes in case of conformational changes. In the sequel, we consider three cases from the *molecular motions DB* [18], see http://www.molmovdb.org/cgi-bin/browse.cgi.

The MSA mode is of special interest since fluctuations directly exploit conformations. It requires a MSA, which we take for granted. For a pair of structures, the nature of conformational changes can be studied using three complementary measures:

- A lRMSD can be computed using the mapping between valid positions provided by the MSA.
- Irrespective of the MSA, a structural alignment can be computed for the two chains. We do so using the structural aligner `Kpax` [19], re-implemented within the `Kpax` package of the SBL.
- Assume domains identified by `SPECTRALDOM` have been obtained. A lRMSD can be computed for each such domain, and these can be combined to provide the *combined RMSD* (RMSD_Comb._) based on the per-domain changes [20]. We use the package Molecular_distances_flexible package from the SBL, see https://sbl.inria.fr/doc/Molecular_distances-user-manual.html.

#### Results: glutamine-binding protein (GlnBP)

GlnBP (226 a.a.) is a protein of ellipsoidal shape consisting of two domains–[21] and Fig. 4. It has been crystallized alone and with its ligand (Gln). Upon binding, GlnBP-Gln exhibits a large-scale movement of the two hinges connecting the two globular domains, with two significant angular changes in the two hinges (41.1 deg. in the Φ angle of Gly89; 34.3 deg. in the *ψ* angle of Glu181) [21]. The two hinges and the two domains are unambiguously detected by `SPECTRALDOM` (Fig. 4), both in the MSA and DM modes (even if this latter mode uses a single structure). The values for the triple (lRMSD, lRMSD for the `Kpax` alignment, RMSD_Comb._) clearly establishes the rigidity of the two domains along this hinge motion (Table 1).

**Table 1:** Fluctuations/MSA mode: structural comparisons for pairs of structures. Columns 3-4: lRMSD computed from the sequence alignment; Columns 5-6: lRMSD computed using the structural aligner `Kpax`; Columns 7-8: combined RMSD RMSD_Comb._ computed from the domains identified by `SPECTRALDOM`. NB: NA values for the (1eps/A, 1g6s/A) owe to the fact that 1eps contains CA only, so that the canonical representation used by `Kpax` (based on *N, C*_*α*_, *C* atoms) cannot be computed.

|  | PDBid/chain | PDBid/chain | #aa | IRMSD | #aa | IRMSD Kpax | #aa | RMSD<sub>Comb.</sub> | k | Fig. |
| --- | --- | --- | --- | --- | --- | --- | --- | --- | --- | --- |
| GBP | 1ggg/A | 1wdn/A | 220 | 5.34 | 164 | 3.27 | 215 | 1.43 | 2 | 4 |
| EF2 | 1n0u/A | 1n0v/C | 818 | 13.60 | 537 | 4.17 | 818 | 4.57 | 2 | 5 |
|  |  |  |  |  |  |  | 818 | 3.90 | 4 | 5 |
|  |  |  |  |  |  |  | 818 | 3.87 | 6 | 5 |

**Figure 4:**
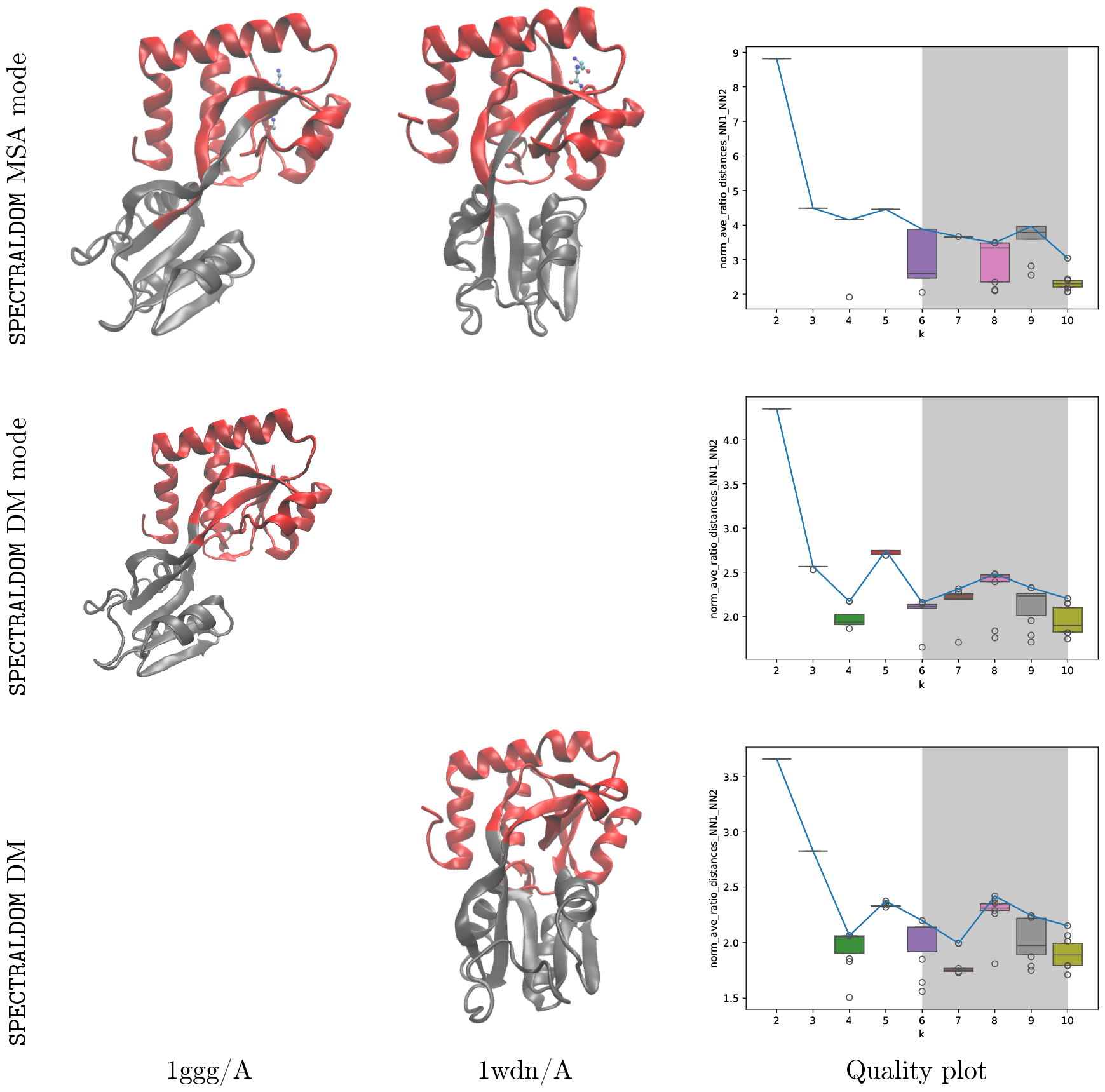
Glutamine-binding protein (GlnBP): comparison of the SPECTRALDOM modes MSA and DM. Crystal structures: unbound: 1ggg/A; bound: 1wdn/A. **(Top)** MSA mode applied to chains A of 1ggg and 1wdn. **(Middle)** DM mode for 1ggg/A. **(Bottom)** DM mode for 1wdn/A.

#### Results: yeast elongation factor (eEF2)

eEF2 (842 a.a.) is a GTP binding protein mediating the translocation of peptidyl-tRNA from the A site to the P site on the ribosome. In yeast, sordarin is a selective inhibitor of eEF2. The conformational changes between unbound and bound (with sordarin) of eEF2 have been studied in detail [22]. To analyse the domains identified, let us recall that six domains are usually considered within eEF2-see [22] and Fig. 5: three N-ter domains: 2-218 or 329-345 (domain I or G-domain), 219-328 (G’ domain), 346-481 (domain II); three C-ter domains: 482-558 (domain III), 559-726 or 801-842 (domain IV), 727-800 (domain V).

**Figure 5:**
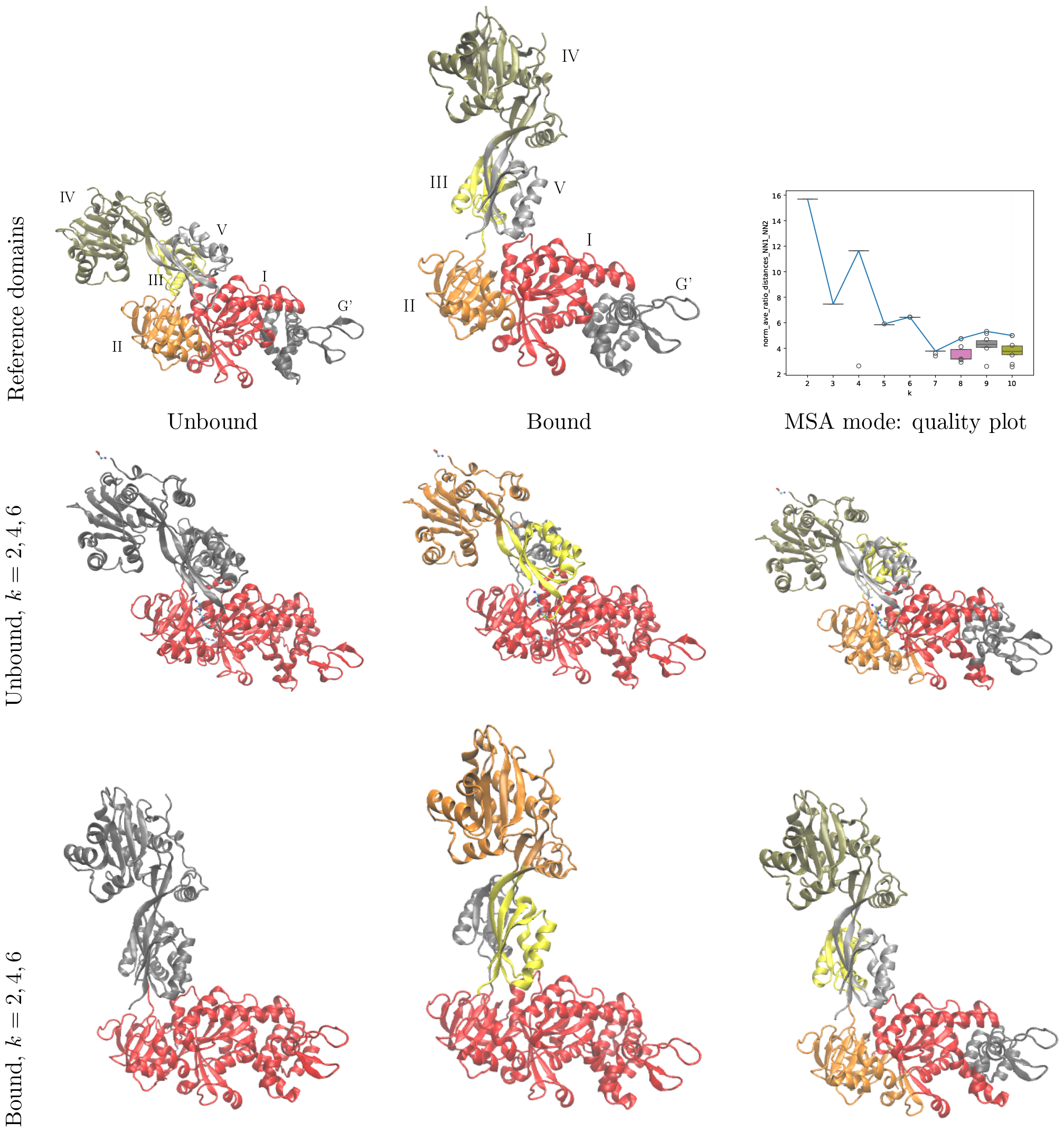
Yeast elongation factor eEF2: analysis with SPECTRALDOM in MSA mode. Crystal structures: unbound: 1n0v/C; bound: 1n0u/A. Values of *k* of interest: *k* = 2, 4, 6 – local maxima with low score variance. **(Top)** Reference domains [22] in for the unbound (1n0v) and bound structures (1n0u), and quality plot for the MSA mode. **(Middle)** Domains of the unbound structure. **(Bottom)** Domains of the bound structure.

The comparison unbound to bound provided by the MSA mode of `SPECTRALDOM` suggests inspecting three decompositions – *k* = 2, 4, 6 (Fig. 5). The first (*k* = 2) shows two meta-domains moving relatively to one another. The three N-ter domains (I, G’, II) are grouped together by `SPECTRALDOM`, which makes sense since they move as a rigid body [22]. The second local maximum (*k* = 4) splits the second meta-domain into the known C-ter domains (III, IV, V). This is coherent with the previous studies, where it is shown along the conformational change unbound to bound, complex motions occur between domains (III, IV, V), and of course with respect to the three N-ter domains. Finally, for *k* = 6, the MSA mode fully recovers the 6 domains as described in the literature. On this example too, the various structural measures indicate that the domains identified are pertinent to study the relative motions at different scales (Table 1).

On such a complicated case, inferring domains from a single structure with the DM mode is more challenging (Fig. 8). The analysis of the unbound case is not very informative. However, that of the bound structure is so. With *k* = 2, the split into three N-ter and C-ter domains is obtained. Also, the last value of *k* with low variance (*k* = 6) identifies the essential features of the six domains.

Altogether, these results show the ability of `SPECTRALDOM` to describe molecular motions at different scales, in particular when exploiting several conformations.

#### Results: the rocking bundle mechanism in LeuT

LeuT has been largely studied as an illustration of transmembrane secondary active transporter: sodium ions flowing down the direction of their gradient provide the energy to transport leucine (and other amino acids) against their gradient. The structure of LeuT presents a couple of intertwining symmetric repeats that deform throughout the cycle in order to transport the substrates. Mechanism-wise, LeuT presents a scaffold domain that supports a bundle of helices translating and rotating during the transport cycle [23].

Since the symmetric repeats do not coincide with the scaffold and bundle regions, LeuT is a prototypical case for distinguishing structural and dynamical domains. Moreover, the extremely low sequence identity between LeuT and other members of its superfamily (0.15 *±* 0.05%) hinders the application of evolutionary-based approaches. The typical compactness of the transmembrane active transporter domains and the complexity of the movement of the bundle make this study case even more challenging.

Among the multitude of structural homologous structures of LeuT, we took PBDid 3tt1 and 3tt3 as examples of the inward-facing (IF) and outward-facing (OF) conformations. `SPECTRALDOM` recovers a maximum for *k* = 4, where one of the four domains (Fig. 6, yellow domain) coincides by 93% with the bundle region. The scaffold region is thus the union of the other three domains: despite this not being the current partition, we notice how different scaffold domains present different displacements (red domain vs. blue and grey domains in Fig. 6), thus providing an explanation for the `SPECTRALDOM` result.

**Figure 6:**
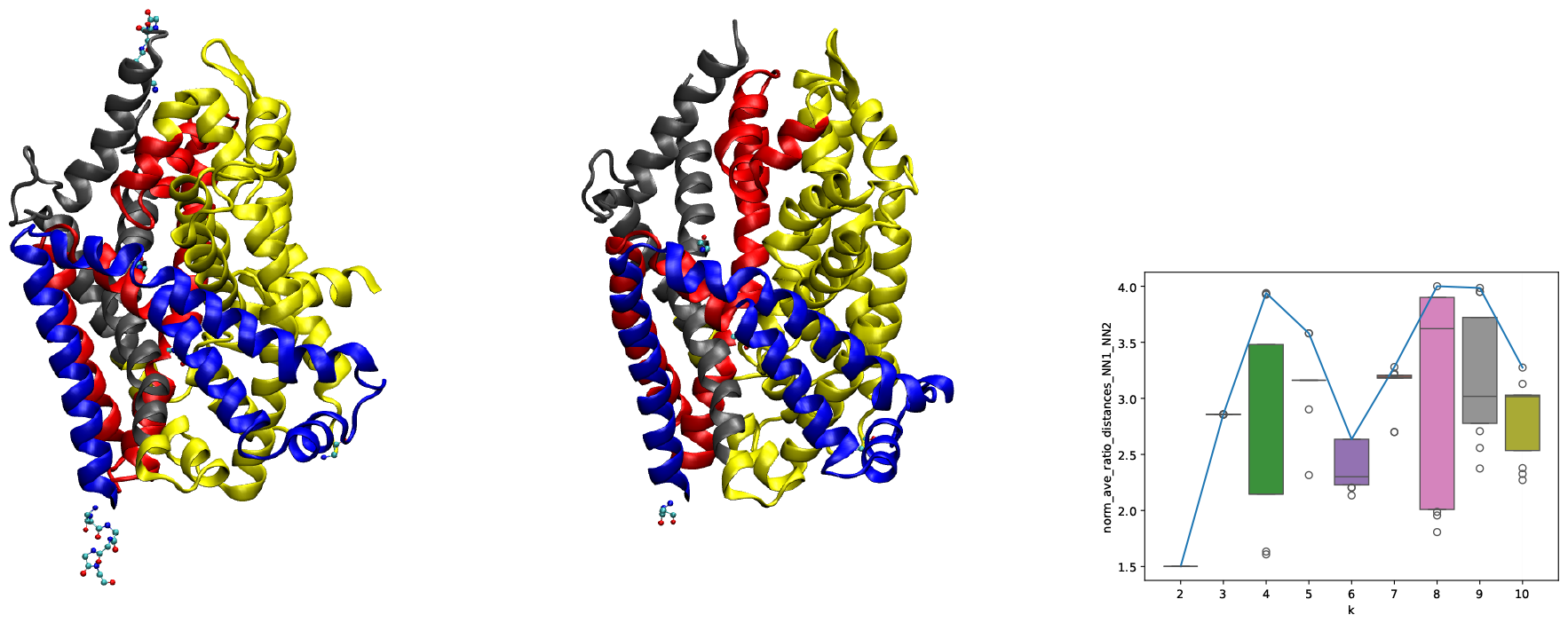
LeuT rocking bundle mechanism: analysis with SPECTRALDOM in MSA mode. Crystal structures: inward-facing: 3tt3/A; outward-facing: 3tt1/A. First value of *k* of interest: *k* = 4. The yellow domain corresponds to the rocking bundle.

On the same target, `Chainsaw` only detects the entire protein as a transmembrane domain, and `SPECTRUS` is not applicable due to the differences in sequence between the two targets. When prompted with only one of the two conformations, it highlights globular domains with low score and no relation with respect to the transport mechanism.

### 3.3 Domains: comparison against a manually curated domain classification

#### Rationale: large-scale domain detection on an evolutionary domain classification

Topology-based structural domain classifications such as CATH and SCOP are often incomplete with respect to homology-based approaches. For this reason, we chose to test our method against `ECOD` [17], whose collection of structural domain has been found to be the most extensive, while remaining consistent with SCOP on the shared domains [24].

#### Results: variation of information on ECOD domains

For each `ECOD` family featuring multi-domain systems, we chose a representative structure and thus collected a set of *∼* 3000 proteins partitioned in their domains. Since `ECOD` domains do not in general cover the whole sequence, for each partitioned protein, we define an extra domain containing all the residues that `ECOD` did not assign to any of its domains.

We then used the *D*-family-matching algorithm again in order to relate `ECOD` and `SPECTRALDOM` domains: each of the two partitions was re-annotated basing on the metacluster indices, then the variation of information VI was calculated on the metacluster partitions. (For *X*_1_ and *X*_2_ two discrete random variables–describing here the assignment of amino acids to domains, denoting *H* the entropy and *I* the mutual information, recall that VI(*X, Y*) = *H*(*X*_1_) + *H*(*X*_2_) − 2*I*(*X*_1_, *X*_2_). VI is null if and only if the two clusterings are equal [25].) The variation of information depends on the number of meta-clusters, but also on the relative values of the number of meta-clusters and the number of `ECOD` domains. For a domain partitioning method (`SPECTRALDOM`, `Chainsaw`), we therefore consider the scatter plot #meta-clusters/#`ECOD` clusters versus VI (Fig. 7). We first note that VI is less that 0.25 for both methods. A one-sided Mann-Whitney U test (a non-parametric two-sample test) to compare the individual values VI_SPECTRALDOM_ versus VI_Chainsaw_ yields a p − value *∼* 10^−11^, showing that the former values are lower than the latter. Remarkably, the partitioning returned by `SPECTRALDOM` is more coherent than that of `Chainsaw` with respect to `ECOD`, without using any evolutionary piece of information.

**Figure 7:**
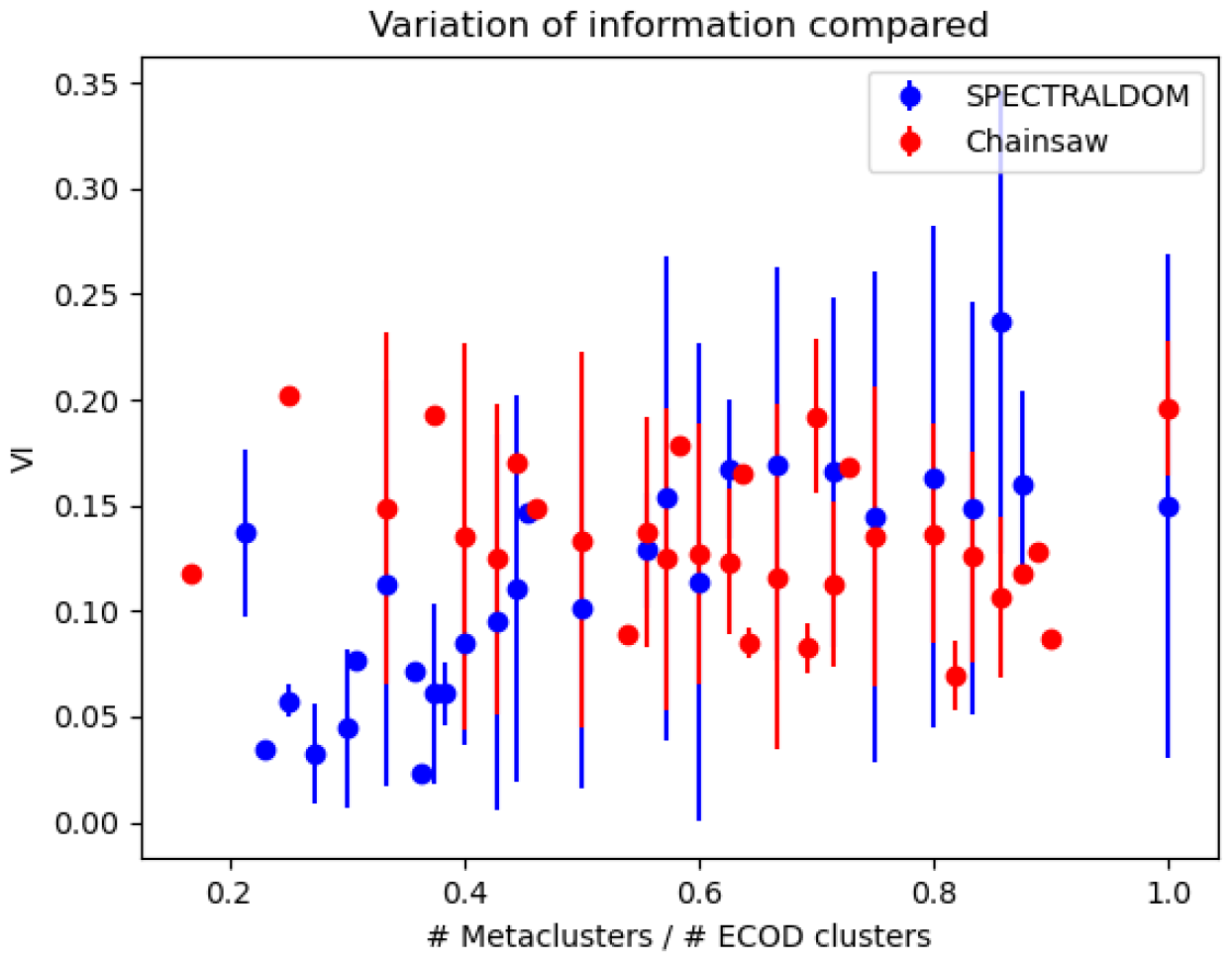
Variation of information (VI) between the clusters of SPECTRALDOM and Chainsaw against those from ECOD. The x-axis reflects the dependency of the VI on the number of labels (metaclusters). When the ratio #metaclusters / #`ECOD` increases, the average VI and its deviation become larger–the coherence with `ECOD` decreases. The one-sided Mann-Whitney U test (a non-parametric two-sample test) shows that the VI values from `SPECTRALDOM` are smaller than those from `Chainsaw`– p − value *∼* 10^−11^.

**Figure 8:**
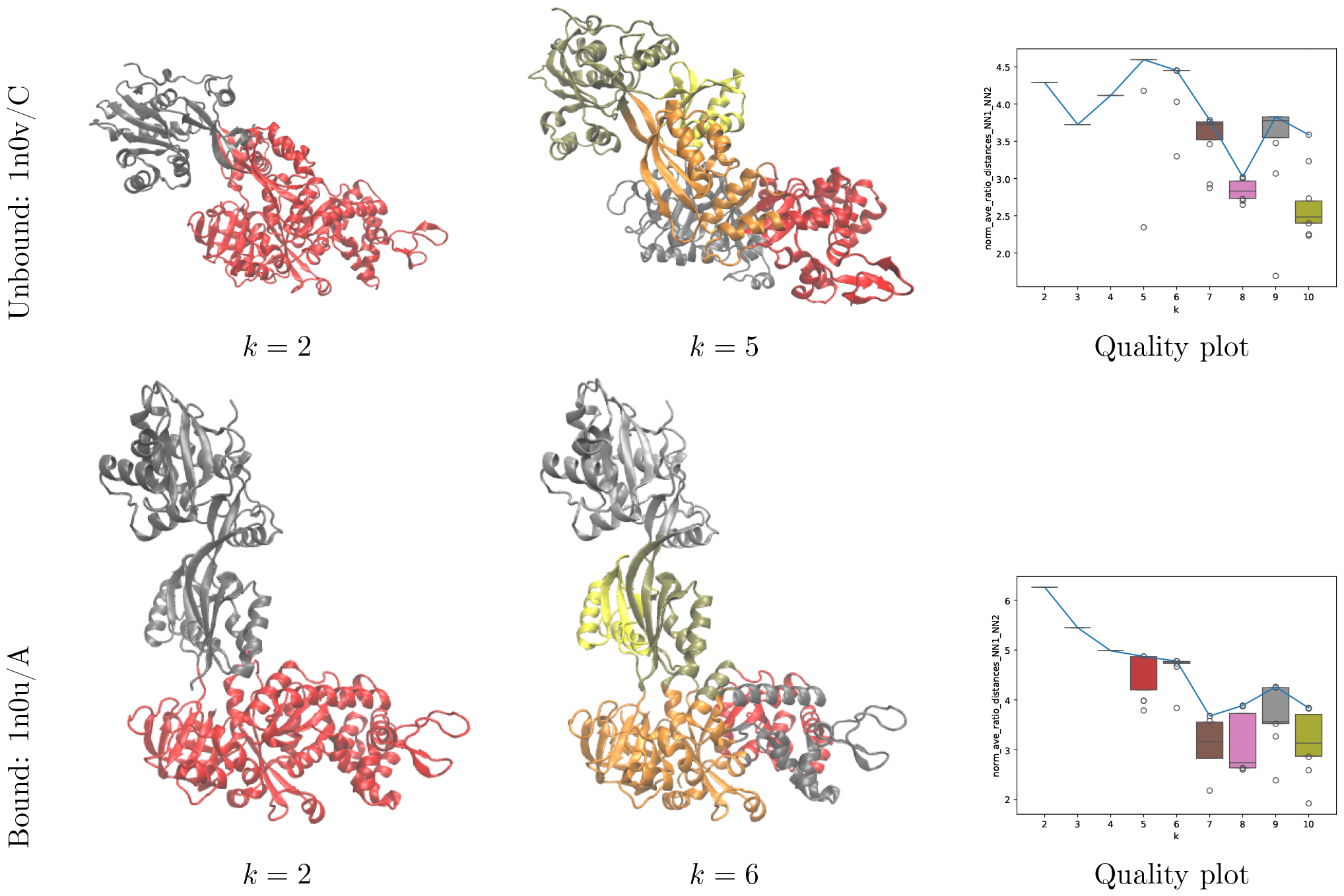
Yeast elongation factor eEF2: analysis with SPECTRALDOM in DM mode.

##### Remark 4

*To assess running times, we used a random subset of 58 structures of the previous dataset. We found* `SPECTRALDOM` *to be on average 5 times faster than* `SPECTRUS`, *and comparable to* `Chainsaw`*– Fig. S4*.

## 4 Outlook

Molecular motions are critical for the activity of biomolecules. They are at play at different scales, ranging from subtle vibrational properties contributing to the free energy of the system, to large amplitude conformational changes repositioning whole structural domains. The automatic identification of such domains is of particular interest as a first step towards the mechanistic explanation of complex mechanisms. Rooted in the vast realm of *graph partitioning / community detection methods*, the original `SPECTRUS` algorithm provides a robust and physically grounded method to identify structural domains. Our work simplifies this method by replacing the computation of normal modes by a simpler *diffusion map* mode relying on a graph Laplacian directly coding pairwise interactions between amino acids. The DM mode significantly reduces the complexity of the method while retaining the physically appealing rationale. We also propose a fully fledged *multiple sequence alignment* mode (MSA), leveraging sequence alignments and fluctuations in experimental structures. The comparison of the DM (single molecule) and MSA (multiple structures) modes stressed the robustness of the method and the coherence of the two modes. Finally, we show how to combat so-called fragmentation issues using a matching between clusters of clusters. Our battery of tests, which handles challenging cases for competing methods, shows that `SPECTRALDOM` compares favorably to contenders while achieving top runtime performances. We anticipate that its integration to the Structural Bioinformatics Library, along with a variety of complementary structure analysis tools, will contribute to its dissemination and broad use.

Two ongoing developments are of particular interest. The first relates to the prediction of mechanisms for macromolecular machines involving a large number of domains (say 10 or more). When these domains move in a rigid fashion with respect to one another, their identification coupled to the ability to sample coherently multiple loop conformations paves the way to deciphering these complex dynamics. This perspective is backed up by our recent experience with resistance-nodulation-cell division (RND) transporters, and membrane proteins in general. The second concerns the ability to identify partially disordered domains, which often occur with deep structure predictors. So far, a direct consequence of the spectral clustering strategy employed in `SPECTRALDOM` is the assignment of all amino acids to domains. Incorporating into `SPECTRALDOM` a disorder predictor and/or a special geometric/topological processing to identify partially disordered regions is a route of choice to handle such cases.

## 5 Artwork

