## Supporting information for "Simpler protein domain identification using spectral clustering"

### S1 Supporting information

Following Section 2.2, `SPECTRALDOM` proposes three modes:

- Mode ENM: the original spectrus with fluctuations (Eq. 1) computed from an elastic network model, and used to compute pairwise similarities.
- Mode DM: stiffness constants are directly used as similarities to derive the symmetric Laplacian used for spectral clustering.
- Mode MSA: fluctuations derived from coordinate changes observed in structures, via a multiple sequence alignment.

**Remark 5** *We also use the MSA mode to handle conformations of a protein. Structures of the same protein obtained by different groups indeed typically exhibit minor changes (a few a.a. added/removed at the N-ter C-ter for example), so that an alignment is needed. Naturally, for conformations come from a simulation, the trivial alignment can be used.*

#### S1.1 Original `SPECTRUS` and its scoring scheme

The original spectrus has been recalled in Section 2.1. In the sequel, we just recall the scoring scheme used to identify plausible values for the number of clusters/domains $k$.

**Score for one run at fixed $k$.** Spectral clustering requires clustering $n$ unit vectors on the sphere $S^{k-1}$. Recall that in `k-means` or k-medoids, one assigns a data point to the nearest center. To assess whether this assignment is stable, one computes the ratio of distances to the first and second nearest centers. (When clusters are well separated, this ratio should be small.) This average ratio for $n$ points is denoted $\overline{r}^{12}_{k,n}$. To compare this value to a null model, let $\overline{r}^{12}_{k,n}[\text{Ref}]$ be the same computed for $n$ random points $S^k$. The `SPECTRUS` fitness score retained for that value of $k$ reads as

$$\text{Score}_{k,n} = \frac{\overline{r}^{12}_{k,n}}{\overline{r}^{12}_{k,n}[\text{Ref}]}. \tag{4}$$

**Best score at fixed $k$.** k-medoids and k-means are randomized algorithms requiring an initial choice of seeds. We use the smart randomized seeding of `k-means++` [15] to get an efficient selection of seeds prior to running `k-means` on $S^{k-1}$. We run `k-means++` $N_r(=10)$ times and retain the partitioning associated with the top score $\text{Score}_{k,n}$.

**Range for $k$ values and repeats.** Consider now a range of values for $k \in [k_{\min}, k_{\max}]$. For each $k$. Running $N_r$ repeats for each value of $k$ yields:

**Definition. 1** *The* quality plot *is the box plot of the score Eq. 4 obtained for $N_r$ repeats for each value of $k \in [k_{min}, k_{max}]$.*

In practice, two files are dumped by `SPECTRALDOM`:

- scores-repeats.csv: the scores obtained for all repeats and all values of $k$.
- scores-best.csv: the best scores for each value of $k$ – thus $N_r$ values in total.

#### S1.2 Sequences, structures, and valid positions

In this section, we explain how the modes ENM, DM and MSA are handled coherently. We assume that MSA are provide in FASTA format. We also assume that MSA are computed from whole primary sequences (seqres entry of the PDB/mmCIF) file, rather than the residues found in the structures.

**MSA mode: sequences, alignments, and (valid) indices, resids.** An *index* is an integer starting at zero. We define two notions of valid indices (Fig. S1).

To make sure that every fluctuation (Eq. 1) is computed with the same number of pairs (that is the binomial coefficient $\binom{\text{num. sequences}}{2}$), we define:

**Definition. 2** *A position is termed* valid *with respect to a MSA provided that there are no dashes (-) in the column associated with this position.*

**Definition. 3** *A position is termed* valid *with respect to a MSA and the associated structures provided that it is valid w.r.t. the MSA and that the corresponding residues are present in their respective structures.*

**Example 1** *Assume one has obtained a MSA for the ectodomain of a protein. Assume that the MSA has length $N$, and that the i-th sequence of each ectodomain has length $n_i < N$ and starts at some offset resid $o_i$:*

- *the MSA focuses only on the a.a. of the ectodomain, that is, the MSA sequence index 0 of the i-th sequence corresponds to the first residue of that sequence in the MSA – and not in the primary sequence.*
- *the MSA sequence index 0 of the i-th sequence corresponds to the residue whose resid is $o_i$.*

The difficulty in manipulating multiple MSA and associated structures resides in indices (Fig. S1):

- alignment indices,
- sequence indices, which are contiguous indices defined from residue sequence numbers (and insertion codes),
- resids found in the structures, which are not contiguous in general due to missing residues.

The main classes are SBL::CSB::T_Multiple_sequence_alignment_DS, SBL::CSB::T_Multiple_sequence_alignment_DS_f and SBL::CSB::T_Chains_residues_contiguous_indexer

Implementation details are provided in the Implementation section of the software, see `https://sbl.inria.fr/doc/Spectral_domain_explorer-user-manual.html`.

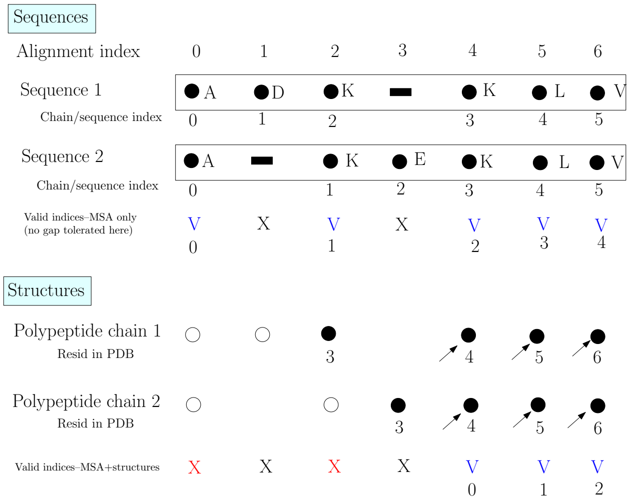


Figure S1: **Sequences, structures, and valid indices.** NB: alignment on sequences are expected to be carried out on primary sequences. **(Top panel: sequences)** Two fictitious sequences with 6 residues, yielding a fictitious MSA with 7 positions. If one does not tolerate any - in a column of the MSA, 5 positions are valid. **(Bottom panel: structures)** The fictitious experimentally resolved structures, with solid (resp. hollow) bullets for residues present (resp. absent) in the structure. On this toy example, it is assumed that the first two residues of each structure are missing. Out of the 5 valid positions, only 3 remain valid if one requires the 2 residues to be present in the structure. These valid residues for the sequence AND the structure have valid indices 0 to 2 for each chain–arrows in the bottom panel.

#### S1.3 ENM and DM modes: the case of a single structure

The previous machinery easily handles the case of a single structure for the ENM and DM modes:

- When loading the structure and its $C_\alpha$ carbons, the corresponding sequence is built – with X letters for residues missed in the structure. The sequence thus has length max_resid.
- A structural filter is used to record missing residues – thus invalid positions in the trivial MSA reduced to one sequence. We use the class SBL::CSB::T_Chains_residues_contiguous_indexer .

#### S1.4 Pseudo-code of SPECTRALDOM

**Algorithm 1 Algorithm `SPECTRALDOM` for a fixed value of $k$.** The pseudo-code handles coherently all modes (ENM, DM, MSA).

```
// 1. Input
1. Load the structures and set the structural filters (one per chain) used in Def. 3
//2. MSA and valid indices
if MSA mode then
   Load the MSA
   Identify valid positions in the MSA and the structures (Def. 3)
else if ENM or DM mode then
   Build a trivial MSA reducing to one sequence
   Build the associated structural filter to identify missing residues (Def. 3)
// 3. Collect CA. In the sequel, they are identified by contiguous indices corresponding to valid positions
in the MSA.
for all chains do
   Collect the C_αs corresponding to valid positions of the MSA
//4. Weights w_ij and associated graph
if ENM mode then
   Identify C_α pairs within a distance threshold
   Compute fluctuation matrix from ENM and the associated weights w_ij
else if DM mode then
   Define the weights w_ij from the stiffnesses – Eq. 3
else if MSA mode then
   For each C_α pair, compute the max distance in all structures
   Identify C_α pairs within a distance threshold
   Compute the fluctuations – Eq. 1
   Compute the associated similarities
Build the weighted graph from similarites
// 5. Spectral clustering
Solve spectral clustering via k-means++ on S^{k-1}
Compute the normalized score Score_{k,n} (Eq. 4)
// 6. Output
Using MSA and valid indices: convert clusters into domains of residues
```

#### S1.5 Managing fragmentation

As a concrete example, we study the case of the `SPECTRUS` analysis of 1cx8 (Fig. S3). The algorithm finds an optimal partition for $k = 3$, but the partition contains several fragments. The partition for $k = 6$, although not relevant per se, is scored as second peak by the `SPECTRUS` quality score, and presents less fragments. Our

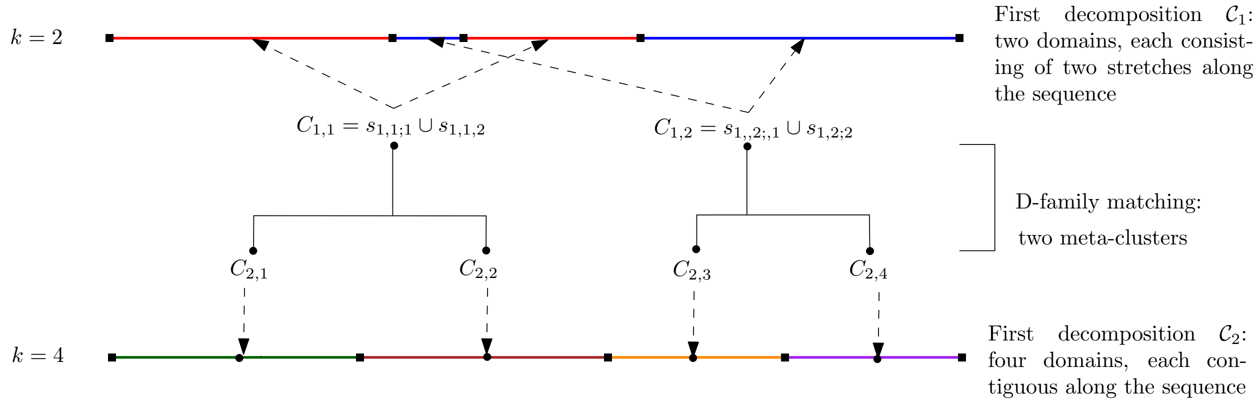


Figure S2: **Reducing fragmentation using the $D$-family-matching algorithm: method.** A chain split into two non-contiguous domains for $k = 2$ (colors: red and blue), and four contiguous domains for $k = 4$ (colors: dark green, brown, orange, purple). The $D$-family-matching algorithm is used to re-assign correctly the short section of the second (blue) domain to the (red) domain $C_{1,1}$. See text for details.

postprocessing procedure is then able to correct the $k = 3$ partition by considering a D-family matching with the $k = 6$ partition. We observe how all fragments that do not persist in the $k = 6$ partition are corrected, and that no additional fragment of the $k = 6$ partition occurs.

#### S1.6 Correlation with `Chainsaw` test cases

We tested `SPECTRALDOM` on all cases reported in [7], and compared the domain partitions using the variation of information VI (Table S1). On the four PDB-deposited structures 1b23, 1a8e, 1jnr, 1cx8, the two methods give comparable results, with a maximum VI of 0.17. It is on the AlphaFold2-predicted structures that the results are much more divergent, with a maximum VI of 0.37.
Indeed, `SPECTRALDOM` always returns a global partitioning where each amino acid is assigned to one cluster, even if it belongs to a region in an unnaturally elongated conformation, which constitutes a typical artifact of structure prediction deep learning models. In order for treating these cases, `SPECTRALDOM` would need an additional preprocessing step filtering all unfolded and elongated regions.

| ID | Chain ID | VI |
|---|---|---|
| 1b23 | P | 0.08 |
| 1a8e | A | 0.03 |
| 1jnr | A | 0.16 |
| 1cx8 | A | 0.17 |
| AF-Q9UJ70 | A | 0.25 |
| AF-P07949 | A | 0.14 |
| AF-Q13002 | A | 0.17 |
| AF-Q15678 | A | 0.37 |

Table S1: **Variation of information between `SPECTRALDOM` and `Chainsaw` partitions over the `Chainsaw` test set.**

#### S1.7 Results: runtimes

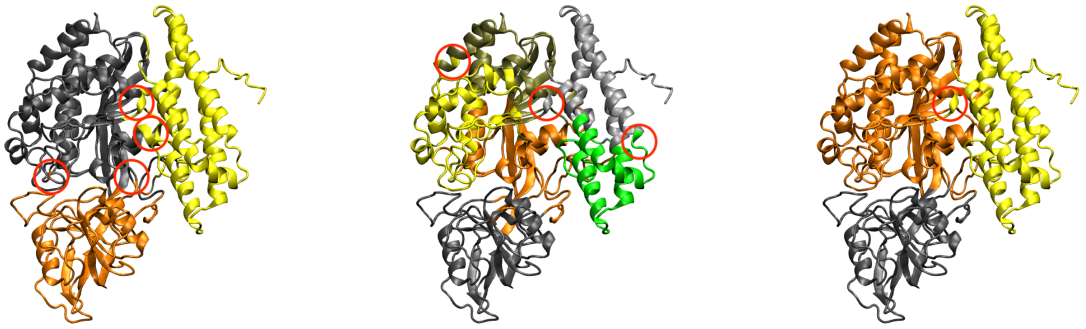

$k = 3$ $k = 6$ $k = 3$ post-processed

Figure S3: **Reducing fragmentation using the $D$-family-matching algorithm: post-processing a `SPECTRUS` case.** PDB 1cx8 partitioned in 3 and 6 clusters. The 3-cluster partition contains multiple fragments (circled), whereas the 6-cluster partition contains less. The latter is thus used to correct the former. Obviously, the fragments that persist in the 6-cluster partition cannot be corrected, such as the helix turn in the middle of the structure.

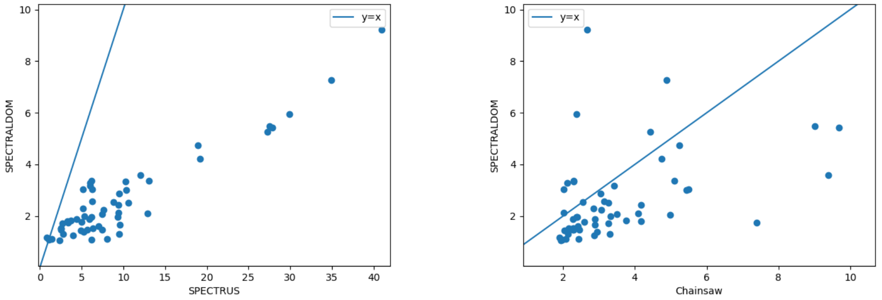


Figure S4: **Runtimes of `SPECTRALDOM` compared to `SPECTRUS` and `Chainsaw`.**
